# Engineering reduced nicotinamide cofactor metabolism for enhanced cell growth and succinic acid production in a succinate dehydrogenase deficient *Yarrowia lipolytica* strain

**DOI:** 10.64898/2026.04.29.721576

**Authors:** Vasiliki Korka, Apostolis Koutinas, Patrick Fickers

## Abstract

**Background:** Succinic acid (SA) is a four-carbon dicarboxylic acid of considerable industrial relevance, with applications spanning the food, chemical, and pharmaceutical sectors. The remarkable acid tolerance of the yeast *Yarrowia lipolytica* makes it a promising microbial cell factory for SA production. Numerous metabolic engineering strategies have focused on disrupting genes encoding the succinate dehydrogenase (SDH) complex to enhance SA accumulation. However, such a modification is associated with impaired growth and the accumulation of by-products, notably acetic acid (AA).

**Results:** To improve growth capacity, SA productivity, and reduce AA formation in *Y. lipolytica* SDH5-deficient strains (*Sdh5Δ*), carbon flux from glycolysis was partially redirected toward the pentose phosphate pathway by overexpression of the native genes encoding glucose-6-phosphate dehydrogenase (*ZWF1)* and 6-phosphogluconate dehydrogenase (*GND1)*, thereby enhancing NADPH generation. The resulting strain was further engineered to increase NADH availability for the mitochondrial electron transport chain by overexpressing genes encoding either a mutated NADPH-dependent malate dehydrogenase (TfMdh) from *Thermus flavus* or the soluble transhydrogenase (EcSthA) from *Escherichia coli*, enabling indirect conversion of NADPH to NADH. This strategy resulted in 2-fold and 2.2-fold increase in SA productivity and titre, respectively, compared to the Sdh5Δ-ALE strain during bioreactor cultivation on glucose-based media. Moreover, AA accumulation was reduced 1.2-fold, while growth rates were significantly improved.

**Conclusions:** The proposed engineering strategies, especially heterologous expression of EcSthA, partly alleviated energy limitations in *Y. lipolytica Sdh5Δ* strain, resulting in improved SA productivity and growth performance.

## Introduction

Succinic acid (SA) is a four-carbon dicarboxylic acid with major industrial relevance and wide range of applications. Current global SA production is approximately 50,000 tonnes per year, of which about 20% is obtained from microbial processes [1]. The SA market is expected to expand significantly, reaching an estimated value of $515.8 million by 2030. The importance of SA relies on its versatility as a precursor for a wide range of high-value chemicals, including adipic acid, 1,4-butanediol, tetrahydrofuran, N-methyl pyrrolidone, 2-pyrrolidone, and γ-butyrolactone. Beyond these applications, SA is widely used in the production of plasticizers, lubricants, food additives, environmentally friendly solvents, pharmaceuticals, and bioplastics [2].

Historically, microbial synthesis of SA has mainly relied on bacterial based bioprocesses. A wide range of wild-type and genetically engineered bacterial strains have been considered, including *Actinobacillus succinogenes* and engineered *Escherichia coli* as well as *Corynebacterium glutamicum* [3]. However, most bacterial hosts require near-neutral pH conditions for optimal growth, under which SA is produced mainly in its dissociated form, succinate. As a result, downstream process recovery for SA involves additional acidification and purification steps, leading to increased process complexity and higher cost. In contrast, yeast species generally display higher tolerance to acidic environments, enabling SA production at pH lower than its pKa values (5.6 and 4.2), greatly simplifying downstream separation and purification, minimizes the need for pH control and the contamination risks [4,5]. The use of cellulose- and starch-rich agro-industrial residues and food wastes has been considered as feedstock to improve the economic feasibility of biobased SA production. Enzymatic hydrolysis of these feedstocks generates glucose-rich hydrolysates, that can be used as carbon and energy sources for cell growth and SA synthesis [2]. Corn steep liquor (CSL) is one of the principal co-products of corn grain refineries that is rich in nutrients, such as organic acids, proteins, amino acids, vitamins, minerals and residual sugars [6]. It has been widely used to produce various organic acids, including citric acid [6,7] and SA by *A. succinogenes* [8].

The yeast *Yarrowia lipolytica* [9] has emerged as an efficient cell factory to produce recombinant proteins, metabolites and ingredients [10,11]. Its exceptional acid resistance makes it particularly well-adapted to produce organic acids, including SA [12,13]. Most engineering strategies for SA accumulation rely on the disruption of the succinate dehydrogenase gene (*SDH*) or attenuation of its activity. While this approach promotes SA accumulation, it also significantly impairs cell growth and carbon source catabolism efficiency. In addition, SA production is often accompanied by the formation of acetic acid (AA) as a major side-product, resulting in the accumulation of acetyl-CoA in the mitochondria. Sdh is an enzyme complex embedded in the inner mitochondrial membrane participating both in the tricarboxylic acid (TCA) cycle by oxidizing succinate to fumarate and in the electron transport chain (ETC) by transferring electron from FADH_2_ to ubiquinone. Although these strategies enable SA accumulation, they also disrupt the TCA cycle, resulting in significantly reduced generation of reduced flavin and nicotinamide cofactors (FADH□ and NADH), thereby diminishing energy supply and cellular growth capacity [2]. To address this, adaptive laboratory evolution (ALE), reconstruction of the reductive TCA cycle (rTCA) in the cytoplasm or mitigating by-product accumulation were successfully used to restore glucose metabolism at least partly, improving cell growth capacity and SA production titre [14–17].

In the present study, we aimed to enhance reduced nicotinamide cofactor generation in a sdh5Δ and sdh5Δ-ALE strains to improve cellular growth capacity and SA productivity. To this end, part of the carbon flux from glucose metabolism was redirected through the pentose phosphate pathway (PPP) by overexpressing the native genes *ZWF1* and *GND1*, encoding glucose-6-phosphate dehydrogenase and 6-phosphogluconate dehydrogenase, respectively. In addition, either a NADPH-dependent malate dehydrogenase from *Thermus flavus* (TfMdh) or the soluble transhydrogenase from *Escherichia coli* (EcSthA) was overexpressed in the ZWF1-GND1 background to indirectly enhance NADH supply to the mitochondrial electron transport chain (ETC) by direct or indirect NADPH conversion.

## Results and discussion

### Strains construction

Auxotroph derivatives of *Y. lipolytica* strains PGC01003 (*Sdh5Δ*, Leu^-^) and PSA02004 (ALE, *Sdh5Δ*, Leu^-^), respectively RIY700 (*Sdh5Δ*, Ura^-^, Leu^-^) and RIY701 (ALE, *Sdh5Δ,* Ura^-^, Leu^-^) served as the starting point of this study (Table 1). Strain PGC01003 was derived from the Po1f (Ura^-^, Leu^-^) laboratory strain by disruption of the *SDH5* gene, which encodes a subunit of the succinate dehydrogenase complex [18]. Although this modification enabled SA accumulation, reaching up to 43 g·L□¹ on crude glycerol, the growth rate of the strain was significantly reduced, particularly on glucose-based media (0.15 ± 0.01 h□¹ on YNBG compared to 0.33 ± 0.01 h□¹ for the prototroph RIY129 strain; data not shown). Glucose metabolism was partially restored through adaptive laboratory evolution (ALE), resulting in strain PSA02004, which exhibited an improved growth rate of 0.17 ± 0.01 h□¹ on YNBG (data not shown). However, its growth rate remained substantially lower than that of the wild-type strain.

**Table 1:**
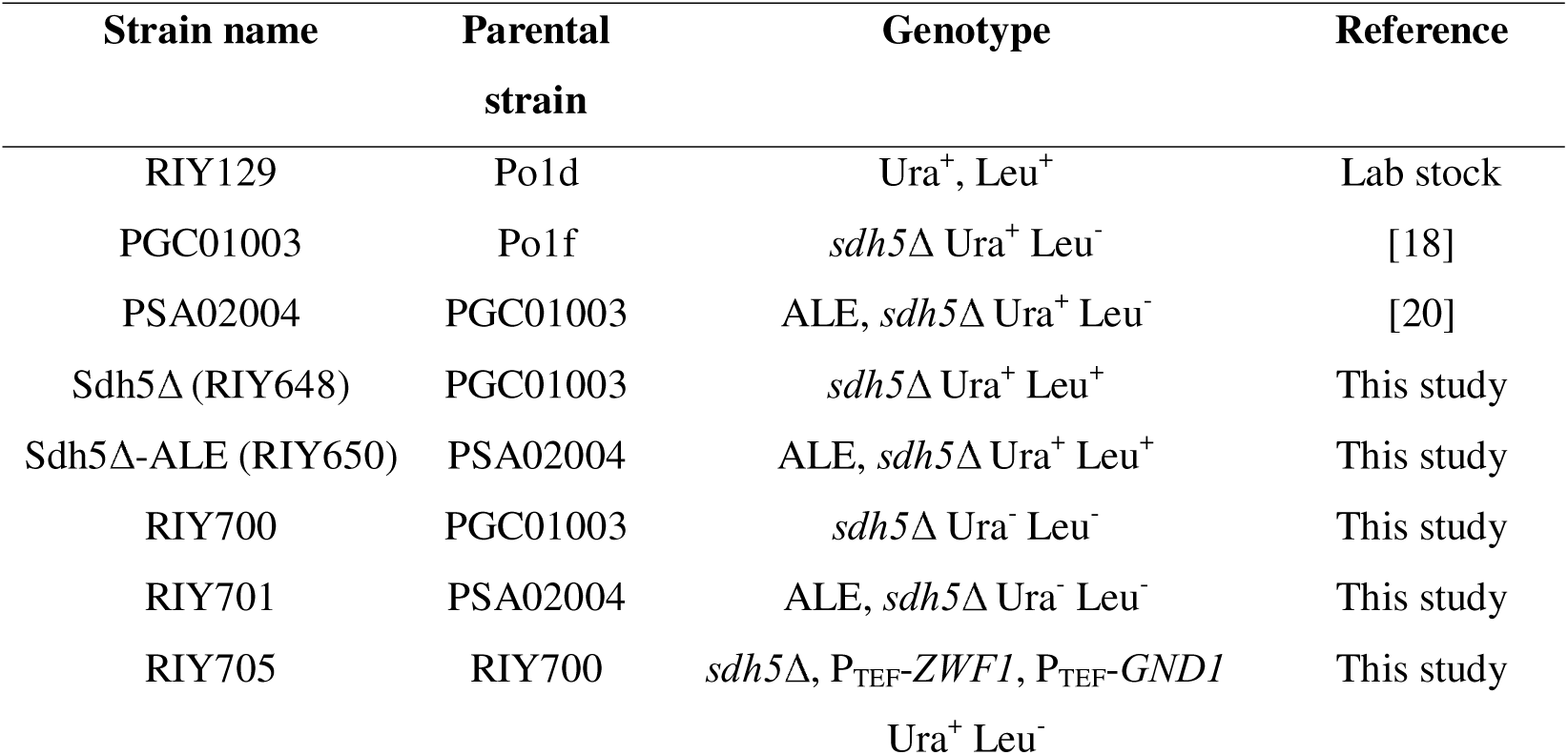
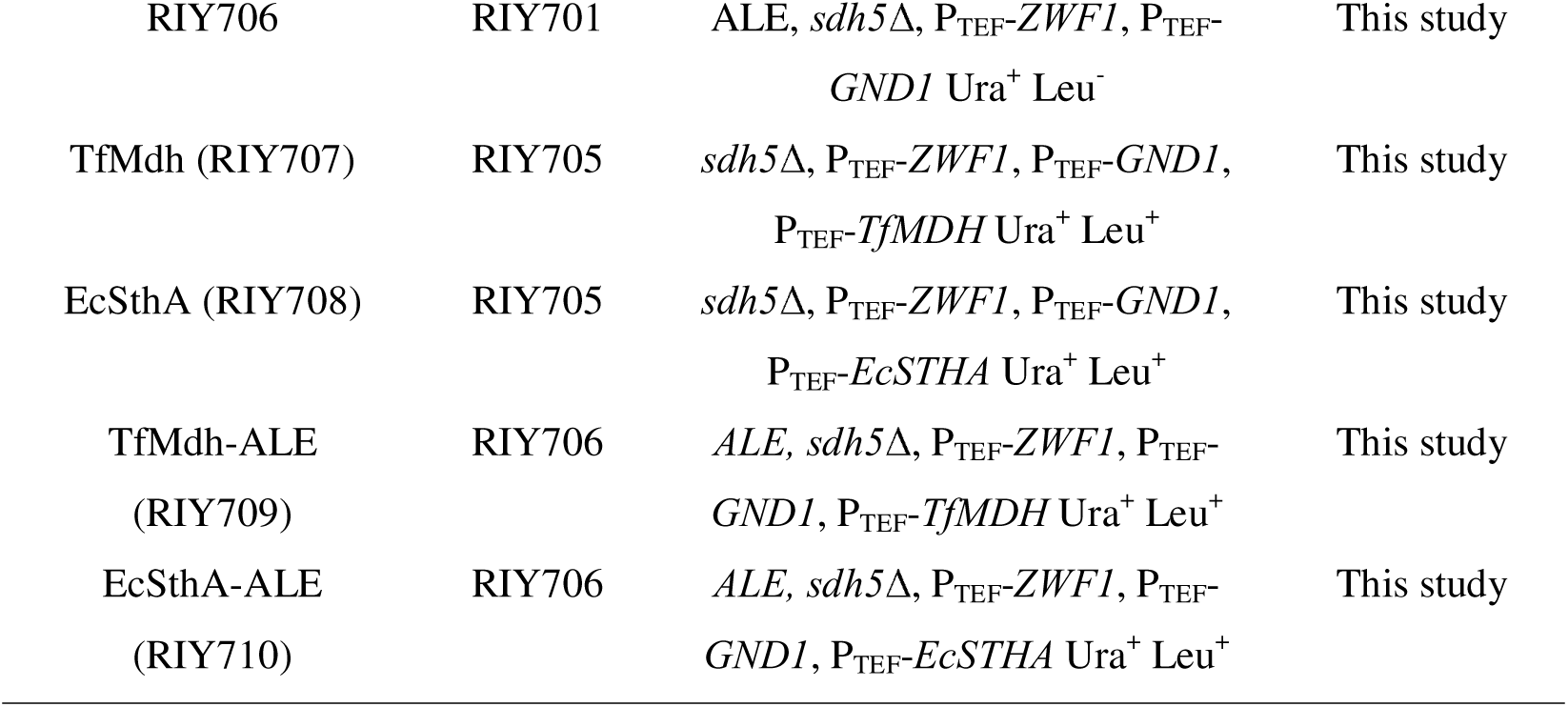
*Y. lipolytica* used in this study.

In this study, we aimed to further enhance the growth capacity of the Sdh5Δ strain. Our strategy involved overexpressing the *ZWF1* (*YALI0E22649g*) and *GND1* (*YALI0B15598g*) genes, which encode glucose-6-phosphate dehydrogenase (G6PDH) and 6-phosphogluconate dehydrogenase (6PGDH), respectively, in the oxidative branch of the pentose phosphate pathway (PPP, Figure 1). These enzymes were selected since they catalyze reactions that generate NADPH from glucose-6-phosphate while producing ribulose-5-phosphate that can subsequently be converted via the non-oxidative PPP into glycolytic intermediates, namely fructose-6-phosphate and glyceraldehyde-3-phosphate (Figure 1).

**Figure 1.**
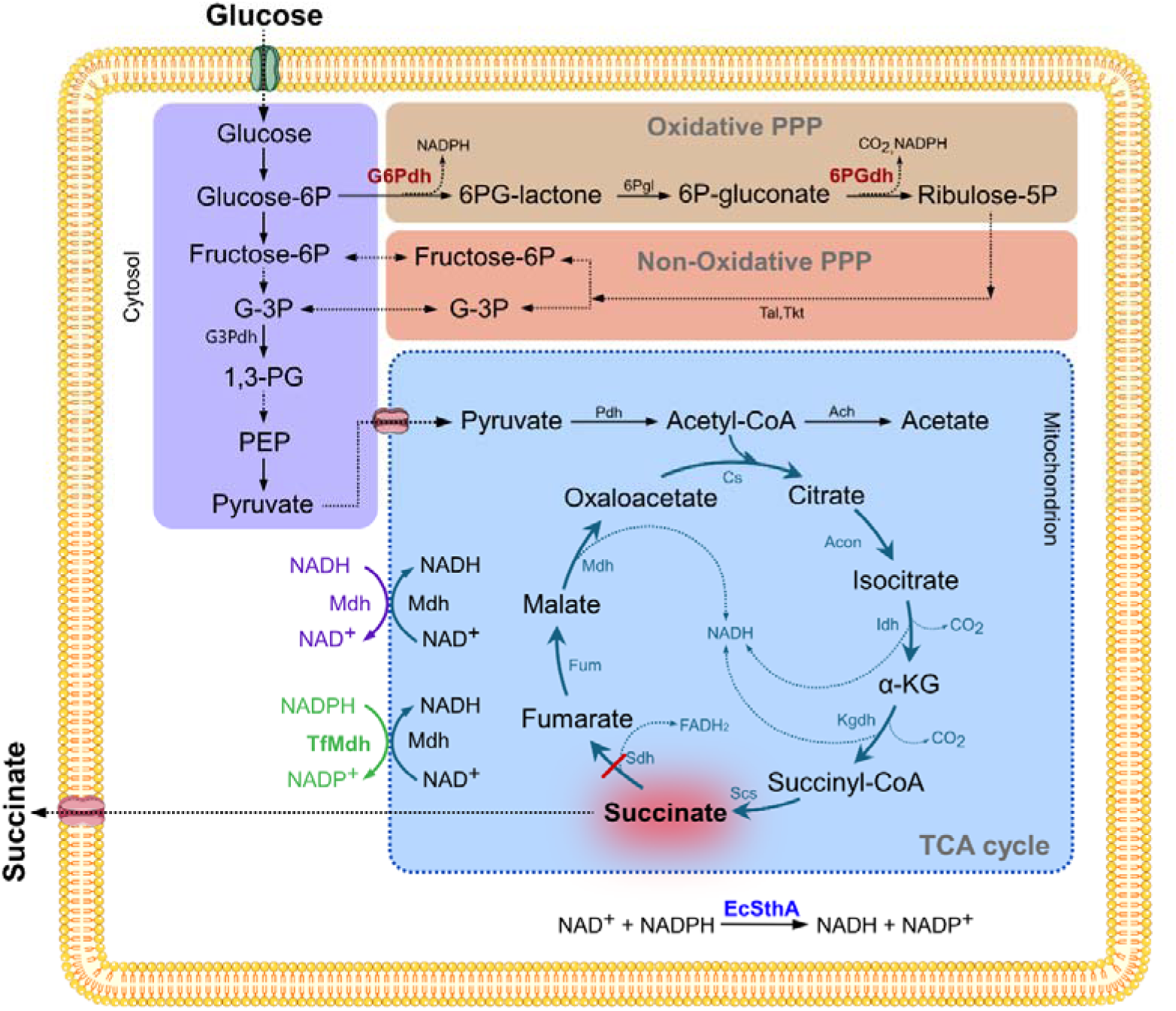
Schematic diagram for succinic acid production by engineered Y. lipolytica strain. Colored boxes represent the glycolysis (purple), oxidative pentose phosphate pathway (brown), non-oxidative pentose phosphate pathway (orange), TCA cycle in the mitochondria with the malate-aspartate shuttle (blue). For malate-aspartate shuttle, only the Mdh/TfMdh and nicotinamide cofactors are shown for simplicity. Dotted arrows indicate multi-step reactions; the red line depicts SDH5 disruption. Genes overexpressed in all constructed strains are shown in red (G6Pdh, glucose-6-phosphate dehydrogenase; 6PGdh, 6-phosphogluconate dehydrogenase), that in the TfMdh and TfMdh-ALE strains in green (TfMdh, NADPH-dependent malate dehydrogenase from T. flavus) and that in EcSthA and EcSthA-ALE strains in blue (EcSthA, soluble transhydrogenase from E. coli). Purple arrows indicate the native Mdh reaction of the malate-aspartate shuttle (MAS), while green arrows indicate the TfMdh reaction in the modified MAS. Enzymes are as follow: Ach, acetyl-CoA hydrolase; Acon, aconitase; Cs, citrate synthase; Fum, fumarase; G-3P, glyceraldehyde 3-phosphate; G3Pdh, glyceraldehyde 3-phosphate dehydrogenase; G6Pdh, glucose-6-phosphate dehydrogenase; Idh, isocitrate dehydrogenase; Kgdh, α-ketoglutarate dehydrogenase; Mdh, malate dehydrogenase; Pdh, pyruvate dehydrogenase; PEP, phosphoenolpyruvate; 1,3-PG, 1,3-bisphosphoglycerate; 6PGdh, 6-phosphogluconate dehydrogenase; 6Pgl, 6-phosphogluconolactonase; Scs, succinylLCoA synthase; Sdh, succinate dehydrogenase; Tal, transaldolase; Tkt, transketolase. Cofactors are as follows: NADPH/NADP, nicotinamide adenine dinucleotide phosphate (reduced/oxidized form); NADH/NAD, nicotinamide adenine dinucleotide (reduced/oxidized form).

To utilize the generated NADPH for energy supply, one approach consisted of expressing the EcSthA gene from *E. coli*, encoding a soluble transhydrogenase, to convert NADPH into NADH in the cytosol. The resulting NADH could then be transported into the mitochondrial matrix by the malate-aspartate shuttle (Figure 1). Alternatively, a mutated TfMdh gene from *T. flavus*, encoding an NADPH-dependent malate dehydrogenase was expressed to enable cytosolic oxaloacetate reduction into malate using NADPH [19]. In both strategies, malate is transported into the mitochondrial matrix by the MAS, where it is reoxidized to oxaloacetate by mitochondrial malate dehydrogenase, generating NADH. This NADH can subsequently feed into the ETC, ultimately contributing to ATP production. In the constructed strains, all genes were overexpressed under the control of the strong constitutive P_TEF_ promoter. Native *ZWF1* and *GND1* genes were expressed in strain RIY700 and RIY701 to give rise to strains RIY705 (*Sdh5Δ, ZWF1, GND1*) and RIY706 (ALE, *Sdh5Δ, ZWF1, GND1*). Codon optimized TfMdh gene was then overexpressed in strain RIY705 and RIY706 to yield strains RIY707 (*TfMdh*, *Sdh5Δ, ZWF1, GND1*; hereafter TfMdh strain) and RIY709 (*TfMdh*, ALE, *Sdh5Δ, ZWF1, GND1*; hereafter TfMdh-ALE). Finally, gene EcSthA was also expressed in strains RIY705 and RIY706 to yield strains RIY708 (*EcSthA*, *Sdh5Δ, ZWF1, GND1*; hereafter EcSthA strain) and RIY710 (*EcSthA*, ALE, *Sdh5Δ, ZWF1, GND1*; hereafter EcSthA-ALE strain), respectively. For each gene, overexpression was verified by RT-qPCR (data not shown).

### Characterization of the constructed strains in shake flasks

As an initial characterization, TfMdh, EcSthA, TfMdh-ALE, EcSthA-ALE and the corresponding parental Sdh5Δ and Sdh5Δ-ALE strains were cultivated in shake flasks in YNBG (glucose 20 g·L^-1^) medium. Biomass, SA, AA and glucose concentrations were determined (Table S1) and the specific production titer for SA (S_SA_) and AA (S_AA_) together with the glucose uptake rate (q_glu_), SA productivity (P_SA_) and yield (Y_SA_) were calculated (Figure 2). As compared to their respective parental strains, EcSthA and EcSthA-ALE strains exhibited a 1.2-fold and 1.5-fold increased biomass while that for TfMdh and TfMdh-ALE were not markedly different (Figure 2A). Compared to Sdh5Δ strain, EcSthA-ALE strain exhibited a 1.7-fold increase in biomass (13.5 vs. 7.9 g_DCW_·L^-1^).

**Figure 2:**
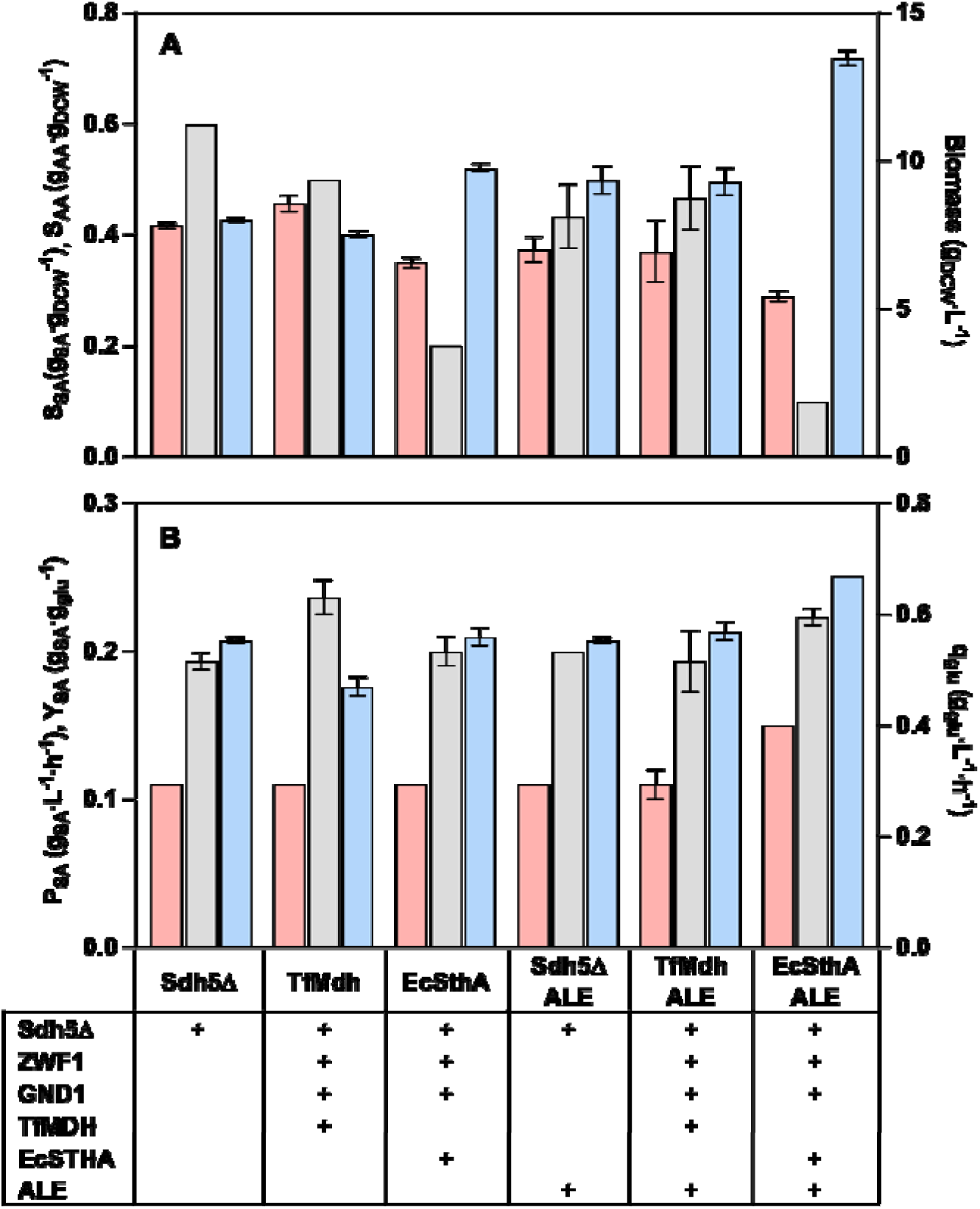
Culture parameters for Sdh5Δ (RIY648), Sdh5Δ-ALE (RIY650), TfMdh (RIY707), EcSthA (RIY708), TfMdh-ALE (RIY709), EcSthA-ALE (RIY710) strains grown in shake flasks in YNBG. Upper panel (A): specific SA titer (S_SA_, red histograms, g_AA_·g^-1^), specific AA titer (S_AA_, grey histograms, g_AA_·g^-1^), biomass (blue histogram, g_DCW_·L^-1^); lower panel (B): SA productivity (P_SA_, red histograms, g**_SA_**·L^-1^·h^-1^); SA yield (Y_SA_, grey histograms, g_SA_·g_glu_^-1^), glucose uptake rate, (q_glu_, blue histograms, g_glu_·L^-1^·h^-1^). Data represents the mean and standard deviation of triplicate cultures.

The remaining glucose concentrations in the medium at the sampling time were different, ranging from 2.3 g_glu_·L^-1^ to 5.7 g_glu_·L^-1^ (Table S1), suggesting differences in the strain performance for glucose catabolism. Importantly, the TfMdh strain displayed a 1.2-fold lower q_glu_ than the Sdh5Δ parental strain (0.47 vs 0.55 g_glu_·L^-1^·h^-1^, respectively) while the EcSthA-ALE strain showed 1.2-fold increased q_glu_ as compared to the Sdh5Δ-ALE strain (0.67 vs 0.55 g_glu_·L^-1^·h^-1^, respectively). In contrast, the values for q_glu_ were not markedly different for EcSthA and TfMdh-ALE as compared to their corresponding parental strain (0.55 vs 0.56 g_glu_·L^-1^·h^-1^and 0.57 vs 0.55 g_glu_·L^-1^·h^-1^, respectively) (Figure 2B). These results highlight that our engineering strategy influence glucose metabolism and cell growth, with effects dependent on the parental strain genotype (ALE vs non-ALE).

**Table S1:** Glucose, biomass, SA and AA concentration during culture of Sdh5Δ (RIY648), Sdh5Δ-ALE (RIY650), TfMdh (RIY707), TfMdh-ALE (RIY709), EcSthA (RIY708), EcSthA-ALE (RIY710) strains grown in shake flasks in YNBG. Data represents the mean of triplicate cultures. Standard deviations were less than 10 % of the average values.

| Strain | Sampling time (h) | Glucose (g<sub>glu</sub>/L) | Biomass (g<sub>DCW</sub>/L) | SA (g<sub>SA</sub>/L) | AA (g<sub>AA</sub>/L) |
| --- | --- | --- | --- | --- | --- |
| Sdh5Δ | 31 | 2.8 | 7.9 | 3.3 | 4.8 |
| TfMdh | 31 | 5.7 | 7.5 | 3.5 | 3.6 |
| EcSthA | 31 | 2.8 | 9.8 | 3.4 | 2.0 |
| Sdh5Δ-ALE | 31 | 2.7 | 9.3 | 3.5 | 4 |
| TfMdh-ALE | 31 | 2.3 | 9.3 | 3.4 | 4.2 |
| ECStHA-ALE | 26 | 2.5 | 13.5 | 4.0 | 2.0 |

In SdhΔ strains, AA is commonly reported as a major side-product [2]. It is derived from acetyl-CoA by acetyl-CoA hydrolase, as acetyl-CoA accumulates in the mitochondria due to disruption of the TCA cycle (Figure 1). Regarding S_SA_ titer, both EcSthA and EcSthA-ALE strains showed reduced S_SA_ as compared to their respective parental strain (1.2 and 1.3-fold, respectively). In contrast, both TfMdh strains showed similar or slightly increase S_SA_ as compared to their parental strain. More marked differences were observed for S_AA_. For the non-evolved strain, both TfMdh and EcSthA showed markedly lower S_AA_ value as compared to the Sdh5Δ strain while for the evolved strain, only EcSthA-ALE showed a markedly reduced S_AA_ value (4.3-fold). Compared to the Sdh5Δ, the reduction is more than 6-fold. In terms of SA productivity, all constructed strain showed values similar to their respective parental strains, except strain EcSthA-ALE with a 1.4-fold increase (0.11 vs 0.15 g_SA_·L^-1^·h^-1^, respectively). Strain TfMdh showed the highest SA yield with value of 0.24 g_SA_·g_glu_^-1^ relative to a Y_SA_ value of 0.19 g_SA_·g_glu_^-1^ for the parental strain.

Overall, EcSthA strains exhibited enhanced growth capacity compared to their respective parental strains, whereas no such improvement was observed in EcMdh strains. In addition, EcSthA strains showed a greater reduction in AA formation, though with lower specific SA production. Notably, the EcSthA-ALE strain displayed an increased glucose uptake rate relative to its parental strain, suggesting a more efficient allocation of glucose toward energy generation. This phenotype is consistent with the overexpression of a heterologous transhydrogenase that enables the conversion of NADPH, derived from the PPP, into NADH to support the mitochondrial ETC. Furthermore, EcSthA-ALE exhibited the highest SA productivity. Relative to its parental strain, the TfMdh strain showed reduced AA production, a slight increase in S_SA_ production, and, most notably, the highest SA yield from glucose. This observation supports the expected role of TfMdh in facilitating the transport and indirect conversion of cytosolic NADPH into mitochondrial NADH via the malate-aspartate shuttle (Figure 1). Based on this initial characterization, the TfMdh and EcSthA-ALE strains were selected for further analysis, as they combined high growth capacity and glucose uptake rates with superior SA productivity and yield.

### Batch cultures in bioreactors with the most efficient mutant strains

As further characterization, TfMdh and EcSthA-ALE strains were grown in 2L batch bioreactor at high glucose concentration (YTG medium) and in a pH-controlled environment. Biomass, glucose concentration, SA and AA titers were determined at various time points over 100 h and compared to that of the Sdh5Δ and Sdh5Δ-ALE parental strains (Figure 3).

**Figure 3.**
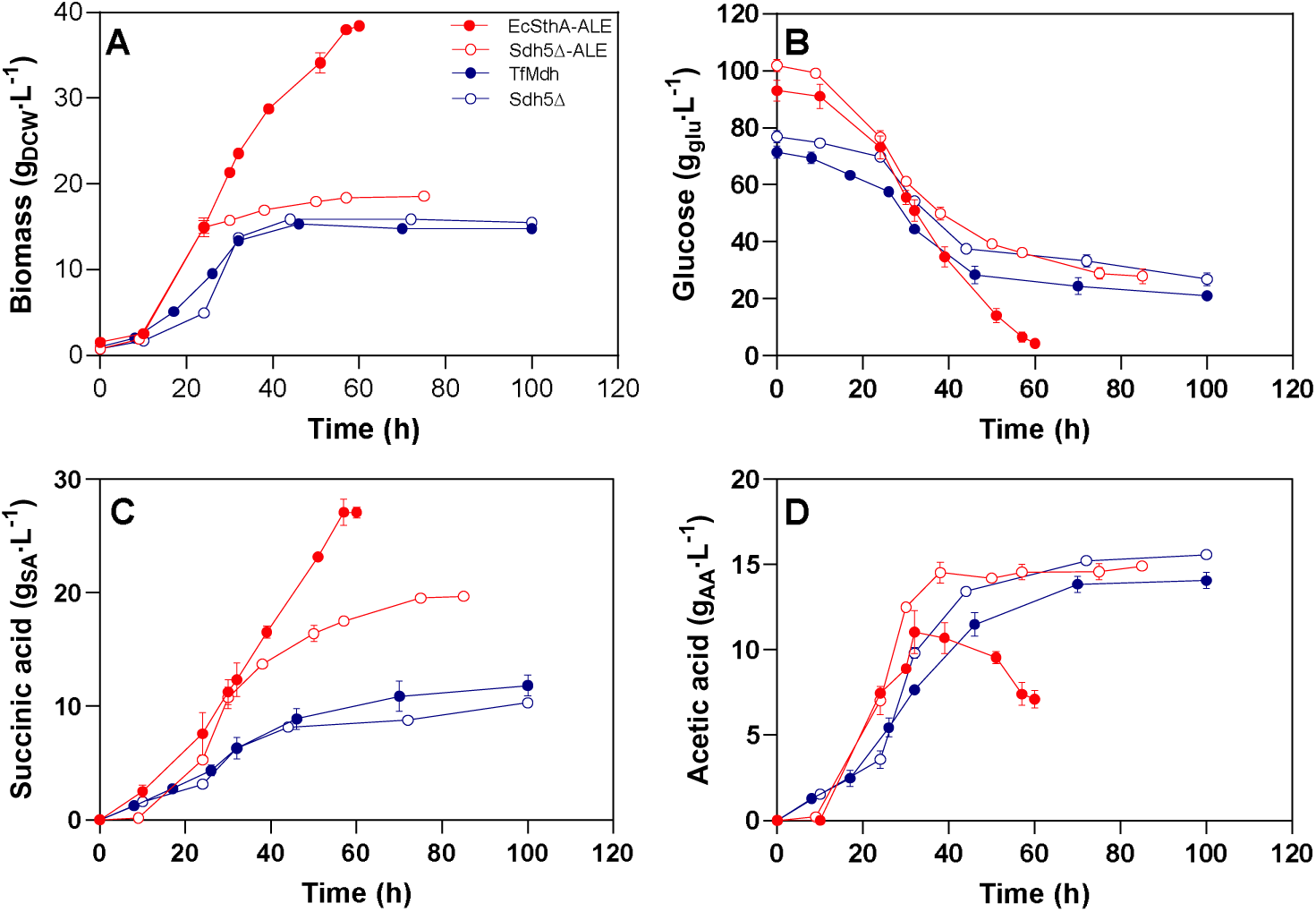
Biomass (A, g_DCW_·L^-1^), glucose (B, g_glu_·L^-1^), succinic acid (C, g_SA_·L^-1^) and acetic acid (D, g_AA_·L^-1^) concentrations during culture in 2L batch bioreactor in YTG medium (glucose 100 g·L^-1^) of EcSthA-ALE (filled red, RIY710), Sdh5Δ-ALE (open red, RIY650), TfMdh (filled blue, RIY707) and Sdh5Δ (open blue, RIY648). Data represents the mean and standard deviation of duplicate cultures.

Similarly, to shake-flask cultures, the EcSthA-ALE strain reached a significantly higher biomass than the parental Sdh5Δ-ALE strain (38.4 vs 18.6 g_DCW_·L□¹, respectively). Although both strains exhibited similar growth during the first 24 h, reaching approximately 15 g_DCW_·L□¹, the growth of the Sdh5Δ-ALE strain was unexpectedly and markedly reduced over the subsequent 51 h (Figure 3A).

Both strains also displayed comparable glucose consumption kinetics during the initial 24 h, with q_glu_ values of 0.83 and 0.99 g_glu_·L□¹·h□¹ for EcSthA-ALE and Sdh5Δ-ALE strains, respectively. However, during the following 36 h, q_glu_ increased to 1.9 g_glu_·L□¹·h□¹ for the EcSthA-ALE strain, while it remained at 1.1 g_glu_·L□¹·h□¹ for the Sdh5Δ-ALE strain. Globally, the glucose uptake rate was 1.7-fold increased for EcSthA-ALE strain as compared to Sdh5Δ-ALE strain, with values of 1.47 and 0.88 g_glu_·L□¹·h□¹, respectively.

The kinetics of SA production followed a similar trend. SA titers reached approximately 11 g_SA_·L□¹ after 30 h of cultivation and further increased to maximal values of 27.1 and 19.6 g_SA_·L□¹ after 60 h and 85 h for EcSthA-ALE and Sdh5Δ-ALE strains, respectively. The SA productivity was respectively equal to 0.45 and 0.26 g**_SA_**·L^-1^·h^-1^.

Regarding AA production, both strains again behaved similarly during the initial phase of cultivation, reaching 11 and 12.5 g_AA_·L□¹ after 32 h for EcSthA-ALE and Sdh5Δ-ALE, respectively. Thereafter, the Sdh5Δ-ALE strain accumulated AA up to 14.4 g_AA_·L□¹ at 38 h, coinciding with growth arrest. In contrast, AA concentration for the EcSthA-ALE strain decreased from 11 to 7.1 g_AA_·L□¹ between 32 and 60 h of cultivation.

In the Sdh5Δ-ALE strain, disruption of the TCA cycle limits the capacity to assimilate acetyl-CoA derived from pyruvate, thereby reducing ATP generation through the ETC. To alleviate this acetyl-CoA overflow, a fraction is converted into acetate via acetyl-CoA hydrolase (Ach). Acetyl-CoA accumulation is known to negatively regulate pyruvate dehydrogenase (PDH) activity, the enzyme that converts pyruvate into acetyl-CoA [21]. Consequently, pyruvate accumulates in the cytosol. In *Pichia kudriavzevii* engineered for L-malic acid production, transcriptomic and metabolomic analyses suggested that pyruvate accumulation contributes to feedback inhibition of glycolysis [22]. Similarly to us, these authors observed a reduced glucose uptake rate after 18 h of growth in rich medium containing 55 g·L□¹ glucose, with approximately 50% of the initial glucose remaining after 48 h of cultivation. Disruption of the TCA cycle also impairs respiration in the ETC, thereby reducing NADH reoxidation. The resulting redox imbalance (elevated NADH/NAD□ ratio) may constrain glycolytic flux by limiting the conversion of glyceraldehyde-3-phosphate to 1,3-bisphosphoglycerate catalyzed by glyceraldehyde-3-phosphate dehydrogenase (Figure 1). This may also explain the observed reduced ability to catabolize glucose over time in the Sdh5Δ and Sdh5Δ-ALE strains as compared to the wild-type strain.

Overexpression of the EcSthA gene in the engineered *P. kudriavzevii* strains restored glucose uptake, resulting in complete glucose consumption withing 48 h and a decreased intracellular pyruvate level by 2.7-fold [22]. Furthermore, expression of EcSthA promoted the interconversion of NADPH and NADP□ in *P. kudriavzevii*, leading to activation of the PPP, as evidenced by the upregulation of *ZWF1* and *GND1* gene expressions, and other related genes. This metabolic rewiring increased the cytosolic NADH supply and was accompanied by elevated transcript levels of PDH. In the EcSthA-ALE strain, expression of the heterologous transhydrogenase gene from *E. coli* likely contributes substantially to the conversion of NADPH into NADH, as supported by the increased q_glu_ value and enhanced growth capacity. The lower AA concentration suggests reduced accumulation of acetyl-CoA within the cell. This improved energy generation in the EcSthA-ALE strain contributes to the significant increase in SA titre.

Although NADPH generation by overexpression of *GND1* and *ZWF1* is expected to be similar in both EcSthA-ALE and TfMdh strains, the indirect conversion of cytosolic NADPH into mitochondrial NADH through the malate-aspartate shuttle mediated by heterologous TfMdh from *T. flavus* appears less efficient than the strategy based on EcSthA transhydrogenase. Importantly, the TfMdh strain did not exhibit higher biomass or SA titers compared to the parental strain, however, it showed a slightly improved glucose uptake rate and reduced AA accumulation. These observations suggest a modest improvement in energy generation in the TfMdh strain and indicate that the heterologous NADPH-dependent malate dehydrogenase is functionally active in *Y. lipolytica*.

### Fed***lJ***batch culture with strain EcSthA-ALE using CSL

In bioreactor-based processes, culture medium costs can represent up to 50% of the total production cost. Consequently, the use of low-cost raw materials or industrial by-products is increasingly considered to reduce overall process expenses. In this context, the EcSthA-ALE strain was cultivated in a fed-batch bioreactor using CSL as an alternative to tryptone and yeast extract, with glucose as the main carbon source. Indeed, CSL has been successfully employed as a low-cost substrate to produce extracellular lipase Lip2p by *Y. lipolytica* in a 2000 L bioreactor [23].

Two media were evaluated; either YTG, containing yeast extract and casein tryptone, and CSLG, in which CSL was used at a concentration adjusted to match the free amino nitrogen (FAN) level of the YTG medium. Cultures were initiated in batch mode with an initial glucose concentration of 100 g·L□¹. When glucose concentration decreased to approximately 50 g·L□¹, feeding was initiated using a concentrated glucose solution (FS, 900 g·L□¹) at different time intervals to maintain a glucose concentration between 20 and 60 g·L□¹ (Figure 4). By the end of the cultivation, the cumulative glucose consumption reached 140 and 125 g_glu_·L□¹ for YTG and CSLG media, respectively.

**Figure 4.**
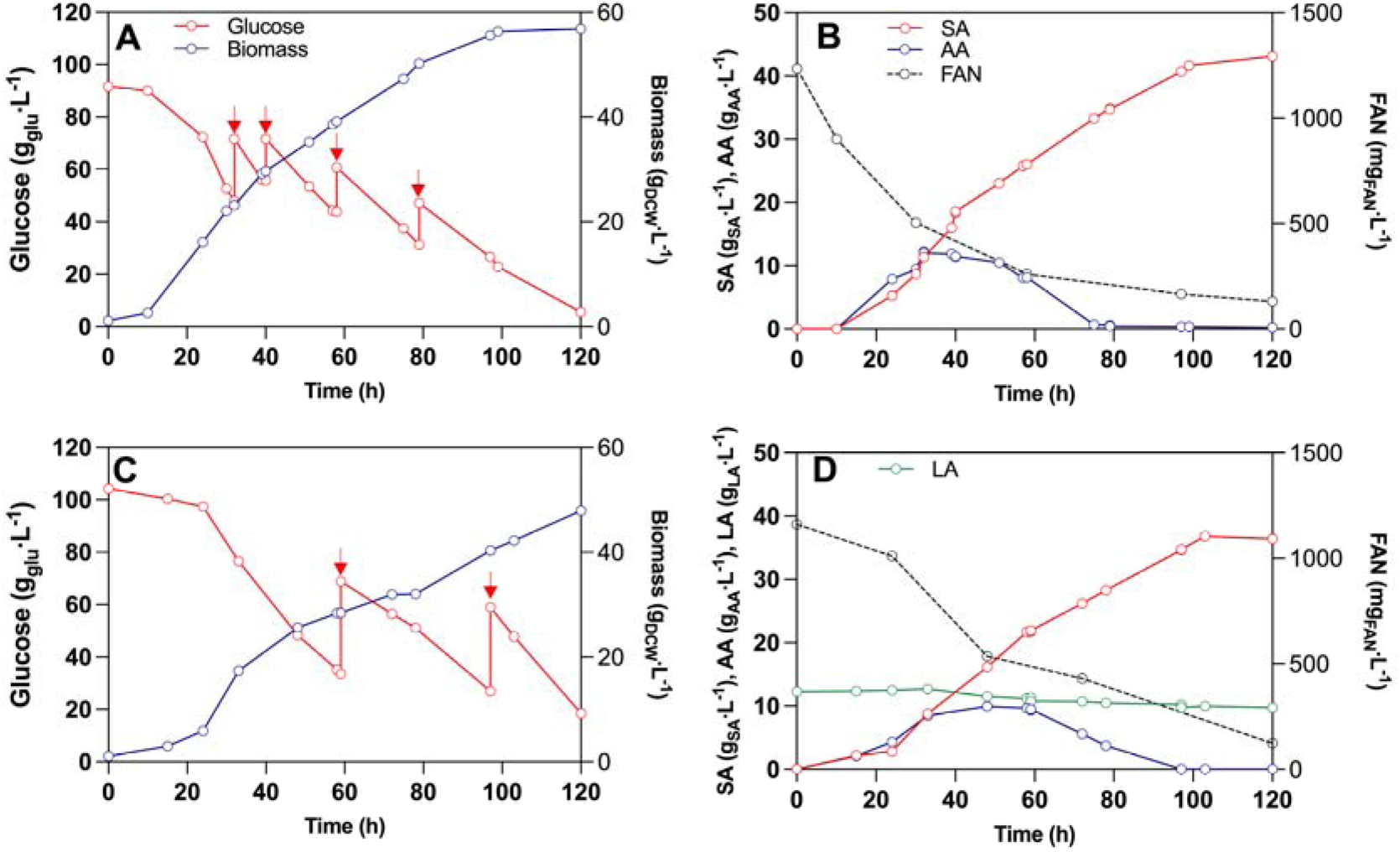
Biomass (blue, panels A and C, g_DCW_·L^-1^), glucose (red, panel A and C, g_glu_·L^-1^), succinic acid (red, panels B and D, g_SA_·L^-1^), acetic acid (blue, panels B and D, g_AA_·L^-1^), FAN (black, panel B and D, mg_FAN_·L^-1^) and lactic acid (green, panel D, g_LA_·L^-1^) concentrations during culture in 6.7 L fed-batch bioreactor in YTG medium (panels A and B) and CSLG medium (panels C and D) of EcSthA-ALE (RIY710) strain. Red arrows indicate FS addition.

In both media, the EcSthA-ALE strain exhibited similar growth profiles, reaching final biomass concentrations of 56 and 48 g_DCW_·L□¹ in YTG and CSLG media, respectively. The slight difference observed may be attributed to variations in glucose availability resulting from the feeding strategy. Importantly, in CSLG medium (Figure 4 C, D), SA production reached 36.8 g_SA_·L□¹ after 103 h, with a yield of 0.30 g_SA_·g_glu_□¹ and a productivity of 0.36 g_SA_·L□¹·h□¹. These values were comparable to those obtained in YTG medium (Figure 4A, B), where SA titers reached 43.1 g_SA_·L□¹, with a yield of 0.31 g_SA_·g_glu_□¹ and a productivity of 0.44 g·L□¹·h□¹.

In both media, AA accumulated similarly, reaching maximal concentrations between 9 and 11 g_AA_·L□¹ after approximately 40 h of cultivation, before being completely reassimilated. This reassimilation occurred slightly faster in YTG than in CSLG medium.

CSL contains lactic acid (LA), with an estimated concentration of approximately 10 g_LA_·L□¹ in CSLG medium. Although *Y. lipolytica* is capable of metabolizing LA (24), it was not consumed under these conditions, most likely due to glucose-mediated catabolite repression. The end of the cultivation coincided with depletion of FAN in both media, indicating that nitrogen limitation was the primary factor restricting further growth and SA production.

Overall, these results demonstrate that the implementation of a fed-batch strategy significantly enhances SA production, and that CSL represents a viable and cost-effective alternative nitrogen source.

## Conclusions

The overexpression of EcSthA gene in Sdh5Δ-ALE strain increased the flux in glucose catabolism as demonstrated by the increased glucose uptake rate in strain in EcSthA-ALE (1.7-fold) in bioreactor batch cultures. This enhanced growth (2.1-fold), SA production (1.4-fold) and productivity (1.7-fold) of the mutant. The fed-batch strategy with the strain EcSthA-ALE further increased SA production reaching up to 43.1 g_SA_·L□¹, with a yield of 0.31 g_SA_·g_glu_□¹ and a productivity of 0.44 g·L□¹·h□¹, while CSL was proved to be an efficient nitrogen source for *Y. lipolytica* growth and SA production.

## Materials and Methods

### Strains, media and culture conditions

The *Y. lipolytica* and *E. coli* strains used in this study are listed in Table 1 and S1, respectively. *E. coli* was grown at 37°C in Luria-Bertani medium (LB), supplemented with antibiotics when required (100 µg·mL□¹ ampicillin or 50 µg/mL kanamycin). *Y. lipolytica* was grown at 28 °C either on YTD (10 g·L□¹ yeast extract, 20 g·L□¹ tryptone, 20 glucose), YTG medium (10 g·L□¹ yeast extract, 20 g·L□¹ tryptone, 100 g·L□¹ glucose), YNBG medium (1.7 g·L□¹ Difco YNB w/o ammonium sulfate and amino acids, 20 g·L□¹ glucose, 5 g·L□¹ NH_4_Cl, 50 mM Na/K phosphate buffer, pH 6.8), YNBGcasa (YNBG with the addition of 2 g·L□¹ Difco casamino acids) or CSLG [175 ml·L□¹ corn steep liquor (CSL) solution, 100 g·L□¹ glucose], the pH was adjusted to 6.8 by addition of NaOH 10M). For CSL solution preparation, 100 mL of raw CSL (Feedstimulans, The Netherlands) was diluted 10-fold with distilled water before being heated at 100 °C for 10 min. After cooling at room temperature, the solution was filtered through Whatman filter paper and autoclaved. Feeding solution (FS) for fed-batch bioreactors was composed of 900 g·L□¹ glucose. For solid media, 15 g·L□¹ agar were added. *Y. lipolytica* transformants were selected on YNB agar plates, supplemented with 0.1 g·L□¹ uracil (YNBG-Ura) or 0.1 g·L□¹ leucine (YNBG-Leu) when necessary to meet auxotrophic requirements or 200 µg·mL□¹ nourseothricin (YTD-Nat).

Yeast cultures were performed at 28°C and 150 rpm in 250 ml shake flasks containing 25 ml of medium or in bioreactor (see below). All cultures were inoculated at an initial OD 600 nm of 0.5 with Na/K phosphate buffer (50 mM, pH 6.8) washed cells from a 12 h preculture in the same medium. For shake flask culture, the pH was initially adjusted to 6.8 and maintained at 6 ± 0.3 by manual addition of 10 M NaOH when requested. Shake flask cultures were performed with three biological replicates. Batch cultures in bioreactor were performed in a 2 L bioreactor (Eppendorf, Bioflo120, Germany) with 1.2 L initial working volume while fed-batch cultures were performed in a 6.7 L bench-top bioreactor (Bioengineering, RALF Advanced) with 3 L initial working volume. They were operated in duplicates at 28°C, with an aeration of 1 vvm and an agitation of 600 rpm. The pH was regulated at 6 with the addition of 10 M NaOH. Carbon and nitrogen sources were sterilized separately from the rest of the medium at 121 °C for 20 min prior to fermentation. For fed-batch culture, FS solution was fed at different time intervals to maintain a glucose concentration in the range of 20-60 g·L□¹.

### General genetic techniques

Standard media and techniques were used for *E. coli* [25]. Restriction enzymes, DNA polymerases, and T4 DNA ligase were obtained from New England Biolabs (NEB) or Thermo Scientific (Thermo Scientific). Primers for PCR and qPCR were synthesized by Eurogentec (Seraing, Belgium, Table S2). Vector TopoBluntII was from Invitrogen. Genomic DNA was purified using a Genomic DNA Purification kit (Thermo Scientific). The vectors used in this study are listed in Table S3. DNA fragments were purified from agarose gels using a NucleoSpin Gel and a PCRclean-up kit (Machery-Nagel). DNA sequencing (Sanger and whole plasmid sequencing) was performed by Eurofin Genomic (Eurofin). Quantitative PCR (qPCR) were performed as described elsewhere with primers listed in Table S2, using the actin genes a reference [26]. Total RNA was extracted using theNucleoSpin RNA Plus kit (Machery-Nagel). qPCR was performed using the Luna Universal qPCR MasterMix and the Step OnePlus Real-Time PCR system (Thermo Scientific). Primers and plasmids design was performed using the software Snapgene (Dotmatics). Gene expression vectors were constructed by Golden Gate (GG) assembly as described elsewhere [27]. *Y. lipolytica* was transformed as described elswhere [28].

**Table S2:**
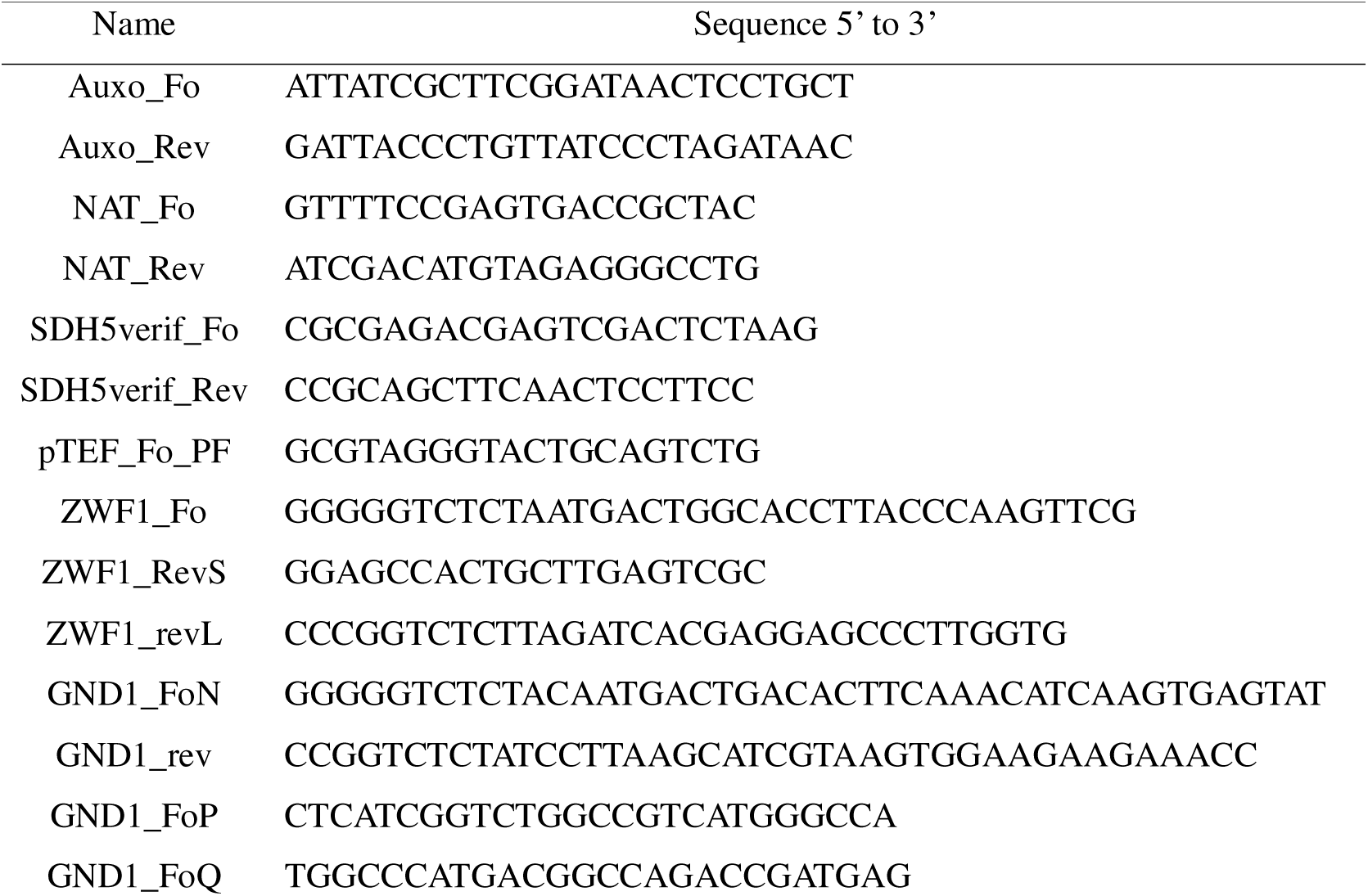
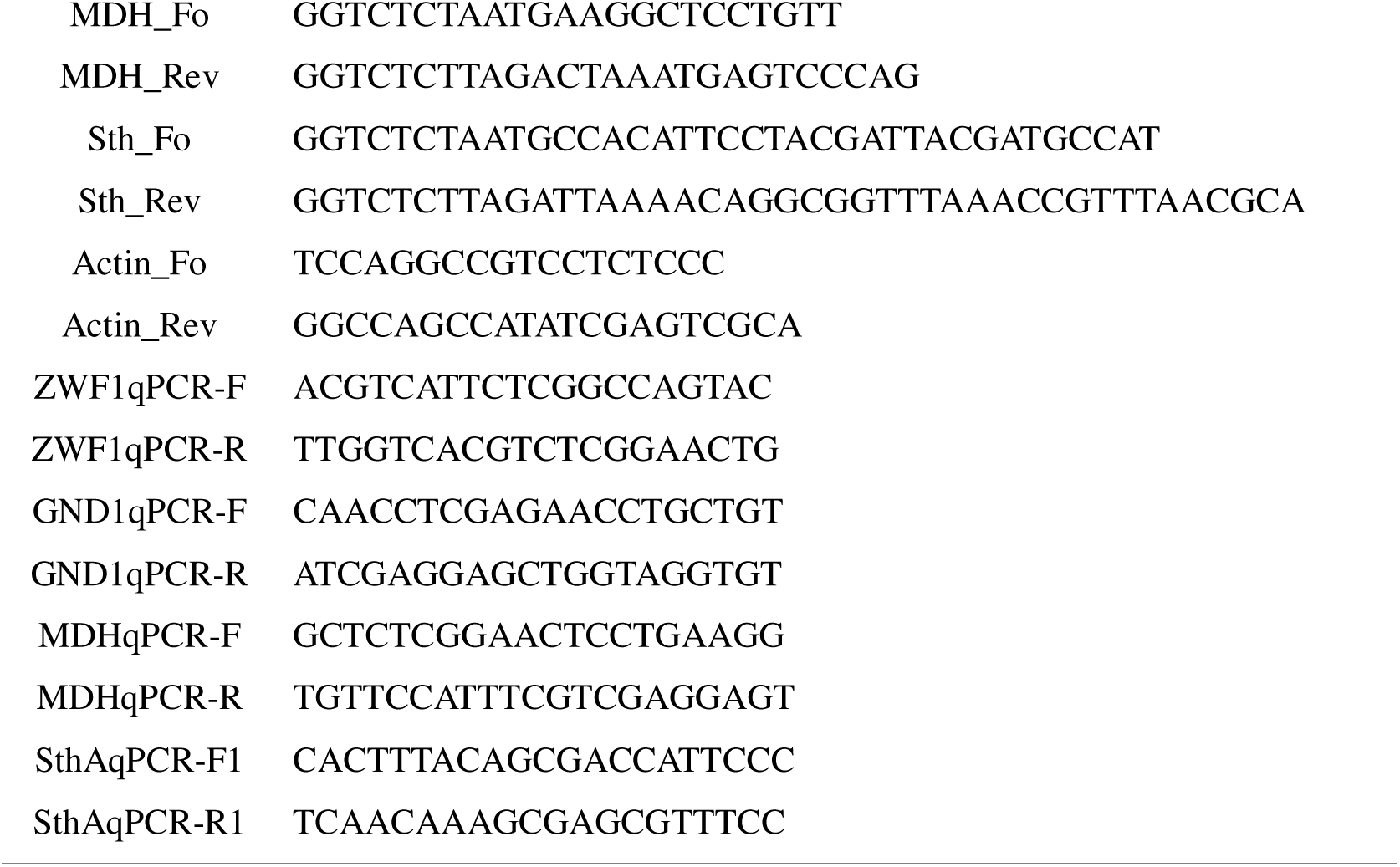
primers used in this study.

**Table S3:** *E. coli* strains and vectors used in this study.

| Strain name (plasmid) | Plasmid-Genotype | Reference |
| --- | --- | --- |
| RIE241 (RIP241) | JMP114, LEU2ex in JMP112 | [29] |
| RIE359 (RIP359) | ARS68 CEN Cre-NAT | Lab stock |
| RIE499 (RIP499) | *GND1* in topo vector | This study |
| RIE500 (RIP500) | *ZWF1* in topo vector | This study |
| RIE501 (RIP501) | P<sub>TEF</sub>-*ZWF1*, P<sub>TEF</sub>-*GND1* | This study |
| RIE502 (RIP502) | TfMDH in topo vector | This study |
| RIE524 (RIP524) | P<sub>TEF</sub>-TfMDH | This study |
| RIE525 (RIP525) | EcSthA in topo vector | This study |
| RIE527 (RIP527) | P<sub>TEF</sub>-EcSTHA | This study |
| A1 (GGE029) | GB3 | [27] |
| A2 (GGE114) | ZetaUP-URA3-RFP-ZetaDOWN | [27] |
| A5 (GGE105) | LEU2 | [27] |
| A9 (GGE085) | P1-P<sub>TEF</sub> | [27] |
| B6 (GGE006) | P2-P<sub>TEF</sub> | [27] |
| D12 (GGE020) | Tlip2 | [27] |
| E1 (GGE021) | Tlip2 | [27] |
| E7 (GGE067) | Zeta-NotI_UP | [27] |
| E8 (GGE038) | Zeta-NotI_DOWN | [27] |

**Figure S1.**
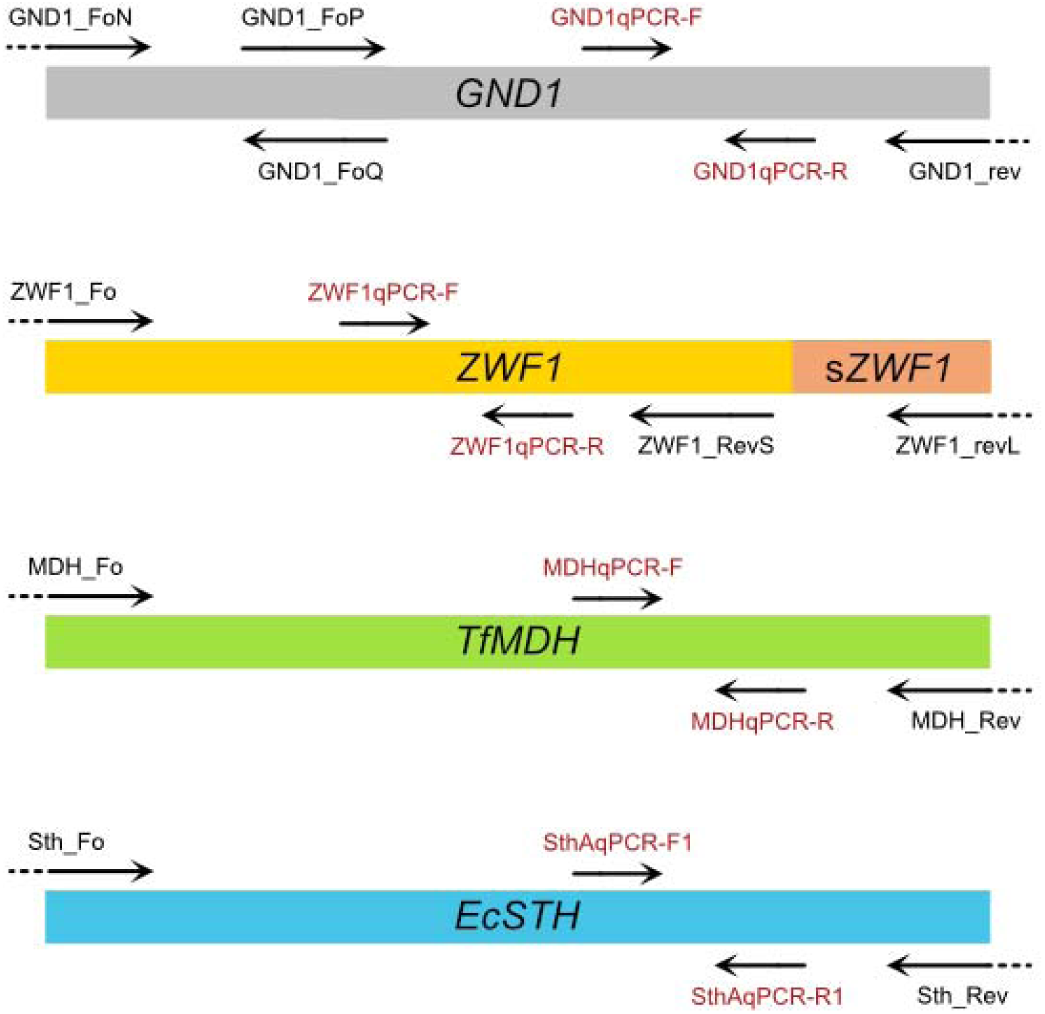
Schematic representation of amplified genes with PCR (black) and qPCR (red) primers; sZWF1 stands for synthetic part of ZWF1 gene. Dots in the arrows show the BsaI sites added at the 5’ and 3’ ends of each gene as stipulated in Materials and Methods.

### Strains RIY648, RIY650, RIY700, and RIY701

Strains PGC01003 (*Sdh5Δ*, Leu^-^) and PSA02004 (ALE, *Sdh5Δ*, Leu^-^) were transformed with *Not*I-digested RIE241 (JMP114). Transformants were selected on YNBG medium, generating prototrophic strains RIY648 (*Sdh5Δ*) and RIY650 (ALE, *Sdh5Δ).* The integration of LEUex marker was verified by PCR using primers Auxo_Fo/Auxo_Rev. In parallel, PGC01003 and PSA02004 were transformed with the replicative vector RIP359 (ARS68 CEN Cre-NAT), and transformants were selected on YTD-Nat. Excision of the URA3ex marker was verified by PCR using primers SDH5verif_Fo/SDH5verif_Rev, yielding auxotrophic strains RIY700 (*Sdh5Δ*, Ura^-^Leu^-^) and RIY701 (ALE, *Sdh5Δ*, Ura^-^ Leu^-^). These strains were subsequently cured of RIP359 by growth on non-selective YTD medium, and vector loss was confirmed by colony PCR using primer NAT_Fo and NAT_Rev.

### Strains RIY705 and RIY706

The *GND1* gene lacking an internal *Bsa*I site was generated by two-step PCR. Fragments of 0.3 kb and 1.9 kb were first amplified from *Y. lipolytica* genomic DNA using primer pairs GND1_FoN/GND1_FoQ and GND1_FoP/GND1_rev, respectively, and fused by overlapping PCR with primers GND1_FoN and GND1_rev to yield a 2.2 kb product (Figure S1). The fragment was cloned into a TopoBluntII vector, generating RIP499 vector. Primers introduced G (ACAA) and H (GGAT) *Bsa*I overhangs [27]. Sequence accuracy was confirmed by DNA sequencing. The *ZWF1* gene was obtained similarly. A 1.8 kb fragment was amplified with primer pair ZWF1_Fo/ZWF1_RevS and fused with a 130 bp ZWF1_end synthetic fragment by overlapping PCR with primer pair ZWF1_Fo/ZWF1_revL (Figure S1). The resulting 1.9 kb product was cloned into a TopoBluntII vector to yield RIP500 vector. *Bsa*I overhangs were introduced as D (AATG) and E (TCTA), and sequence accuracy was confirmed.

Vector RIP501 was assembled by Golden Gate using *ZWF1* and *GND1* as G1 and G2 modules, respectively, with parts GGE114, GGE085, RIP500, GGE014, GGE006, RIP499, and GGE0021 (Table S3). Assembly was verified by sequencing. *Not*I-digested RIP501 was used to transform *Y. lipolytica* strains RIY700 and RIY701, yielding RIY705 (*Sdh5Δ*, *ZWF1*, *GND1*) and RIY706 (ALE, *Sdh5Δ*, *ZWF1*, *GND1*) strains. Transformants were selected on YNBGcasa, and integration of P_TEF_-ZWF1 and P_TEF_-GND1 cassettes was confirmed by PCR with primer pairs pTEF_Fo_PF/ZWF1qPCR-R and pTEF_Fo_PF/GND1qPCR-R, respectively. Overexpression of the *ZWF1* and *GND1* genes in strains RIY705 and RIY706, compared to their respective parental strains, was verified by qPCR using the primer pairs ZWF1qPCR-F/ZWF1qPCR-R and GND1qPCR-F/GND1qPCR-R, respectively.

### Strains RIY707, RIY709, RIY708 and RIY710

The protein sequence of the tMDH-EX7 mutated malate dehydrogenase from *Thermus flavus* [19] was back translated into DNA and codon optimized using the SnapGene software and the *Y. lipolytica* CLIB22 codon usage table (https://www.kazusa.or.jp). The resulting DNA fragment containing D (overhang AATG) and E (overhang TCTA) type *Bsa*I restriction site at 5’ and 3’ ends was synthesized (Twist bioscience) and named RIP502. Vector RIP524 was assembled by GG using parts GGE029, GGE067, GGE105, GGE085, RIP502, GGE020, and GGE038 (Table S3), and verified by sequencing. *Not*I-digested RIP524 was used to transform RIY705 and RIY706 strains, yielding RIY707 (*TfMdh*, *Sdh5Δ*, *ZWF1*, *GND1*) and RIY709 (*TfMdh*, ALE, *Sdh5Δ*, *ZWF1*, *GND1*) strains. Integration of the PTEF-TfMDH-EX7 cassette was confirmed by PCR with primer pairs pTEF_Fo_PF/MDHqPCR-R, and expression was verified by qPCR with primers pairs MDHqPCR-F / MDHqPCR-R. The SthA gene from *E. coli* DH5α was amplified by PCR with primer pairs STH_Fo/STH_Rev, cloned into TopoBluntII vector, and the resulting RIP525 vector was sequence-verified. Vector RIP527 was constructed by Golden Gate as above, replacing RIP502 with RIP525 and sequence verified. *Not*I-digested RIP527 vector was used to transform RIY705 and RIY706 strains, yielding RIY708 (*EcSthA*, *Sdh5Δ*, *ZWF1*, *GND1*) and RIY710 (*EcSthA*, ALE, *Sdh5Δ*, *ZWF1*, *GND1*) strains. Integration of the PTEF-EcSthA cassette was confirmed by PCR with primer pair pTEF_Fo_PF/SthAqPCR-R, and expression was verified by qPCR with primer SthAqPCR-F / SthAqPCR-R.

## Analytical methods

Cell growth was monitored either by optical density at 660nm (OD600) using a spectrophotometer (U-2000 Hitachi) or by dry cell weight (DCW). An OD600 value of 1 was found to correspond to 1.42 g_DCW_·L□¹. Cells were collected by centrifugation and washed twice with deionized water. The cells were dried at 60 °C for 48 h and then the sample was weighted. Sugars and organic acids were determined using a Shimadzu HPLC system with a Shimadzu RI detector and a Rezex ROA-Organic acid H+ column. The temperature of the column was set at 65 ◦C and the mobile phase was 10 mM H_2_SO_4_ aqueous solution at 0.6 ml·min□¹ flow rate. Monosaccharides were also determined with a Shodex SP0810 (8.0 × 300 mm) column. The temperature of the column was 80 °C and the mobile phase was HPLC grade water at a flow rate of 1.0 ml·min□¹. Free amino nitrogen (FAN) was determined according to Lie, 1973 [29].

## Calculation of cultivation kinetic parameters

Specific growth rate (h^-1^) was calculated as:

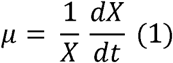

where *X* is the biomass (g_DCW_·L□¹) and *t* is the culture time (h).

Specific S_SA_ titer (g_SA_·g_DCW_□¹) was calculated as:

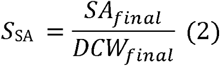

With SA_final_ and DCW_final_ the biomass dry cell weight and the concentration of succinic acid in g·L□¹ at the end of the culture.

Specific S_AA_ titer (g_AA_·g_DCW_□¹) was calculated as:

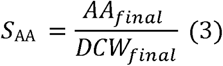

With AA_final_ the concentration of acetic acid in g·L□¹ at the end of the culture.

SA yields (g_SA_·g_glu_□¹) for batch (4) and fed batch (5) cultures were calculated as:

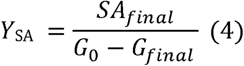

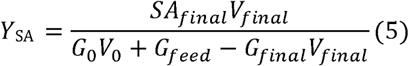

With V_0_ and V_final_ the volume in L at the beginning and at the end of the culture. G_0_, and G_final_ is the glucose concentration in g·L□¹ in the bioreactor at the beginning and at the end of the culture. G_feed_ is the glucose fed in the bioreactor in g.

SA productivity (g_SA_·L□^1^·h□¹), was calculated as:

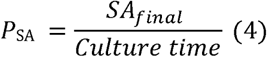

With SA_final_ the concentration in SA in g·L□¹ at the end of the culture (h). Glucose consumption rate (q_glu_·L□^1^·h□¹) was calculated as:

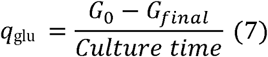

## Acknowledgements

The authors wish to thank Prof. Carol Lin from School of Energy and Environment at City University of Hong-Kong for providing *Y. lipolytica* strain PGC01003 and PSA02004.

## Author contributions

PF and VK designed the experiments and analyzed the results. VK performed the experiments and prepared the draft of the manuscript. PF and AK secure the fundings. All the authors contributed to editing the draft and approved the final manuscript.

